# FluoVolt Staining Induces Photodamage During Live-Cell Voltage Imaging

**DOI:** 10.64898/2026.06.08.730801

**Authors:** Evren M. Akyuz, Manuela I. Mitroi, Fleur Groualle, Foteini Patera, Ruman Rahman, Stuart J. Smith, Ian Spendlove, Judith M. Ramage, Hester Franks, Andrew M. Jackson, Adam M. Blanchard, Anna A. Malecka, Frankie J. Rawson

**Affiliations:** School of Veterinary Medicine and Science, University of Nottingham, Sutton Bonington, UK; Biodiscovery Institute, University of Nottingham, Nottingham, UK; Centre for Cancer Sciences, School of Medicine, University of Nottingham, Nottingham, UK; School of Pharmacy, University of Nottingham, Nottingham, UK; School of Pharmacy, University of Lincoln, Lincoln, UK; School of Life Sciences, University of Nottingham, Nottingham, UK; Deprtment of Oncology, Nottingham University Hospitals NHS Trust, Nottingham, UK

## Abstract

Fluorescent voltage-sensitive dyes (VSDs) enable non-invasive, high-throughput optical measurement of membrane potential in living cells, but the analytical reliability of such measurements depends critically on whether the dye and associated imaging conditions perturb the system under study. Here, we systematically characterise the photophysical performance and cell-perturbing effects of FluoVolt, a widely adopted VSD, across cancer cell lines (GIN31 glioblastoma and SK-MEL-30 melanoma) and primary human macrophages. Photobleaching kinetics were strongly cell-type-dependent, with SK-MEL-30 cells exhibiting complete fluorescence loss within 400 seconds under standard widefield conditions. FluoVolt staining combined with laser excitation caused an approximately 2.5-fold increase in cell detachment relative to unstained controls, and dual-wavelength excitation (488 + 405 nm) reduced GIN31 cell viability by approximately 17.5%. Critically, morphological changes, a transition from elongated to amoeboid-like phenotypes, were detected under staining conditions alone, prior to any laser exposure, indicating baseline dye-induced perturbation independent of phototoxicity. Halving dye concentration and loading time significantly attenuated these effects while preserving measurable fluorescence signal. These findings identify FluoVolt staining and excitation as previously uncharacterised sources of systematic measurement artefact and provide practical, actionable guidance for protocol design, control selection, and data interpretation in optical membrane potential imaging.

## Introduction

Membrane potential (V_mem_) is a fundamental biophysical property of all cells, arising from the unequal distribution of ions across the plasma membrane and maintained by ion channels, transporters, and pumps^1,2^. This electrochemical gradient is essential for numerous physiological processes in both excitable cells, such as neurones and muscle cells, and non-excitable cells, including epithelial and immune cells^3–6^. Beyond its classical role in electrical signalling, V_mem_ has been implicated in regulating cell cycle progression, migration, differentiation, and metabolism^7–10^. In addition to this, V_mem_ is frequently dysregulated in cancer cells, driving cell proliferation and migration among other effects^9,10^. In immune cells, such as macrophages (Mφs), V_mem_ dynamics are closely linked to activation state, polarisation, and inflammatory responses, highlighting a broader role for bioelectric signalling in immune regulation^11,12^.

The current gold standard for measuring V_mem_ is patch-clamp electrophysiology, in which a glass micropipette forms a high-resistance seal with the cell membrane to directly record ionic currents and voltage changes^13,14^. While highly sensitive and quantitative, this technique is inherently low throughput, technically demanding, and limited to measurements in a small number of cells^13,15^. Furthermore, patch-clamp recordings are typically localised, providing measurements at the single-cell level and therefore being less suited for investigating dynamic activity across large cell networks^16^.

To overcome these limitations, fluorescent voltage-sensitive dyes (VSDs) have been developed to enable optical measurements of V_mem_. These probes allow non-invasive, high-throughput imaging of voltage dynamics across many cells simultaneously^17,18^. These dyes can be broadly classified into fast-response electrochromic probes and slower redistribution-based probes^17,19^. While fast-response dyes offer high temporal resolution allowing V_mem_ recordings in the millisecond range, slow-response dyes are often more suitable for steady-state or long-term measurements due to their increased signal stability^17,19,20^.

Genetically encoded voltage indicators have also been developed as an alternative approach for optical monitoring of V_mem_. These transgenes are engineered into cells to express fluorescent protein-based sensors that enable real-time optical detection of changes in V_mem_, offering the advantage of cell-type-specific targeting^21^. However, their application may be limited by lower signal intensity, slower response kinetics, and more complex implementation, particularly in primary cells^21,22^. Consequently, small-molecule dyes remain widely used^22^.

FluoVolt is a commercially available VSD designed to provide rapid and sensitive detection of V_mem_ changes in living cells and enables detection of rapid voltage fluctuations while also being designed to support measurements over longer imaging periods^23^. It operates via a photoinduced electron transfer (PET) mechanism, in which a fluorophore is linked to an electron-rich donor group^23,24^. Upon excitation, fluorescence from the fluorophore can be quenched by electron transfer from the donor to the excited fluorophore. The rate of this process is influenced by the electric field across the cell membrane; changes in V_mem_ alter the efficiency of PET, thereby modulating the degree of fluorescence quenching^23,24^. As a result, variations in V_mem_ produce corresponding changes in fluorescence intensity^23,24^. FluoVolt has been widely adopted due to its compatibility with standard fluorescence microscopy and its ability to capture dynamic changes in V_mem_ without the need for invasive electrophysiological techniques.

To support robust investigation of bioelectric signalling in cancer cell models (glioblastoma and melanoma cell lines) and human monocyte-derived Mφs (hMDMs), we aimed to establish optimised conditions for optical V_mem_ measurements using FluoVolt. As reliable optical recordings depend on both stable fluorescence signals and preservation of cellular health, we systematically assessed how key imaging parameters influenced fluorescence performance and cellular integrity across the different cell types. Parameters such as imaging duration and excitation intensity were assessed to determine their effects on fluorescence signal stability and photobleaching. We also monitored changes in cellular integrity during imaging, including alterations in cell morphology, viability, and cell detachment from the culture surface. These experiments were performed to identify imaging conditions that allowed optical membrane potential measurements to be carried out while minimising adverse effects on the cells.

## Materials and Methods

### Cell culture

GIN31 glioblastoma cells were previously isolated from the 5-aminolevulinic acid (5-ALA)-positive invasive margin of a glioblastoma tumour at Queen’s Medical Centre, Nottingham^25,26^. Cells were authenticated by short tandem repeat profiling (Eurofins) and cultured in Dulbecco’s modified Eagle medium (DMEM; Gibco) supplemented with 10% foetal bovine serum (FBS; Gibco).

SK-MEL-30 melanoma cells were authenticated by short tandem repeat profiling (Eurofins) and cultured in Roswell Park Memorial Institute 1640 medium (RPMI 1640; Sigma-Aldrich) supplemented with 10% FBS (Gibco).

All cells were maintained at 37 °C in a humidified atmosphere containing 5% CO₂ and were routinely tested for mycoplasma contamination.

### Ethics

Ethics approval for studies involving GIN31 cells was obtained from the East Midlands-Derby Research Ethics Committee (REC Reference: 25/EM/0160). All procedures were conducted in accordance with relevant guidelines and regulations.

Studies involving hMDMs were conducted under ethics approval granted by the University of Nottingham Research Ethics Committee (Reference: NB-161-1711). Each donor provided informed consent and signed a written consent form before blood donation and only the blood of healthy donors was used in the study.

### Monocyte isolation and macrophage differentiation

Peripheral blood was obtained from healthy donors via venepuncture using syringes loaded with heparin (Wockhardt). Whole blood was diluted with Dulbecco’s phosphate-buffered saline (DPBS; Sigma-Aldrich) and layered onto Lymphoprep (Serumwerk Bernburg AG) without mixing. Samples were centrifuged at 400 × g for 25 min at 21 °C with low deceleration.

The peripheral blood mononuclear cell (PBMC) layer was collected and washed three times with DPBS, followed by one wash in cold MACS buffer (DPBS supplemented with 1% FBS and 2 mM ethylenediaminetetraacetic acid (EDTA)). CD14⁺ monocytes were isolated using magnetic microbeads (Miltenyi Biotec) as per protocol.

Cell purity was assessed by flow cytometry using a fluorescein isothiocyanate (FITC)-conjugated anti-CD14 antibody (Miltenyi Biotec) on a MACSQuant Analyzer 10 (Miltenyi Biotec) and analysed using FlowJo software (Waters Biosciences). Standard purity was >95%.

Purified monocytes were cultured in RPMI 1640 (Sigma-Aldrich) supplemented with 10% FBS (Sigma-Aldrich) in ultra-low attachment plates (Corning Costar) at densities of 5 × 10⁶ cells per well (6-well, 5 mL) or 1 × 10⁶ cells per well (24-well, 1 mL) for 5 days.

For differentiation, granulocyte-macrophage colony-stimulating factor (GM-CSF; PeproTech) was added at 20 U/mL to generate pro-inflammatory ‘M1’ Mφs, and macrophage colony-stimulating factor (M-CSF; ImmunoTools) was added at 10 ng/mL to generate anti-inflammatory ‘M2’ Mφs. Differentiation supplements were added at the time of plating and fresh medium containing cytokines was added on day 4.

Mφs were harvested on day 5 by incubation on ice for 20 minutes in cold DPBS followed by gentle pipetting, washed, and replated in tissue culture-treated plates (Thermo Scientific Nunclon). Typical yields ranged from 40% to 80% of the initial monocyte input.

### FluoVolt staining and imaging

V_mem_ was assessed using the FluoVolt membrane potential kit (Invitrogen, Thermo Fisher Scientific). Cells were seeded and allowed to adhere for 24 hours prior to experimentation. Cells were washed twice with phosphate-buffered saline (PBS) prior to staining.

The loading solution was prepared in PBS containing FluoVolt dye (1:1000) and PowerLoad concentrate (1:100). Cells were incubated for 20 minutes at room temperature, followed by two washes with PBS. Cells were imaged in PBS unless stated otherwise.

Fluorescence imaging was performed using a Nikon Eclipse Ti2 equipped for widefield microscopy with GFP (488nm) and DAPI (405nm) filter sets. All imaging parameters were kept constant across conditions.

Image analysis was performed using Fiji (ImageJ). Fluorescence intensity was quantified on a per-cell basis using region-of-interest measurements and corrected for background signal.

### Fluorescence time-course analysis and curve fitting

FluoVolt fluorescence intensity was quantified over a 10-minute imaging period for each condition. For each cell type, measurements were obtained from three independent biological replicates and averaged at each time point.

The averaged fluorescence intensity data were analysed using GraphPad Prism. Nonlinear regression was performed using a four-parameter logistic (4PL) model to fit sigmoidal curves to the time-course data. The fitted curves were used to interpolate fluorescence intensity values, enabling comparison and visualisation of multiple cell types on a single graph.

### Cell viability

Cell viability was determined using the Trypan Blue exclusion assay. Following imaging and/or laser exposure, the supernatant containing suspended cells was collected and mixed 1:1 (v/v) with Trypan Blue solution (Sigma-Aldrich).

The mixture was loaded onto a haemocytometer, and cells were counted manually under a light microscope. Viable (unstained) and non-viable (stained) cells were distinguished based on dye exclusion.

Cell viability (%) was calculated as the number of viable cells divided by the total number of cells, multiplied by 100.

### Cell morphology analysis

Cell morphology was quantified by measuring the aspect ratio (length-to-width ratio) of individual cells. Microscopy images were analysed using Fiji (ImageJ).

Images were calibrated using the scale bar to obtain measurements in micrometres (µm). Cell length was defined as the longest axis of the cell, and cell width was measured perpendicular to this axis at the widest point. The aspect ratio was calculated as the ratio of cell length to cell width.

Cells were classified based on aspect ratio, with values > 3 defined as elongated, values < 2 defined as amoeboid-like, and values between 2 and 3 defined as intermediate.

A minimum of 25 cells per condition were analysed across 3 independent images, with the same 3 fields of view maintained across wells to account for intra-well variability. Cells that were overlapping, out of focus, or partially within the field of view were excluded from analysis.

### Statistical analysis

All statistical analyses were performed using GraphPad Prism.

For fluorescence time-course experiments (Figure 1), data from three independent biological replicates were averaged and fitted using nonlinear regression with a 4PL model, as described above. No statistical comparisons were performed for these data.

**Figure 1:**
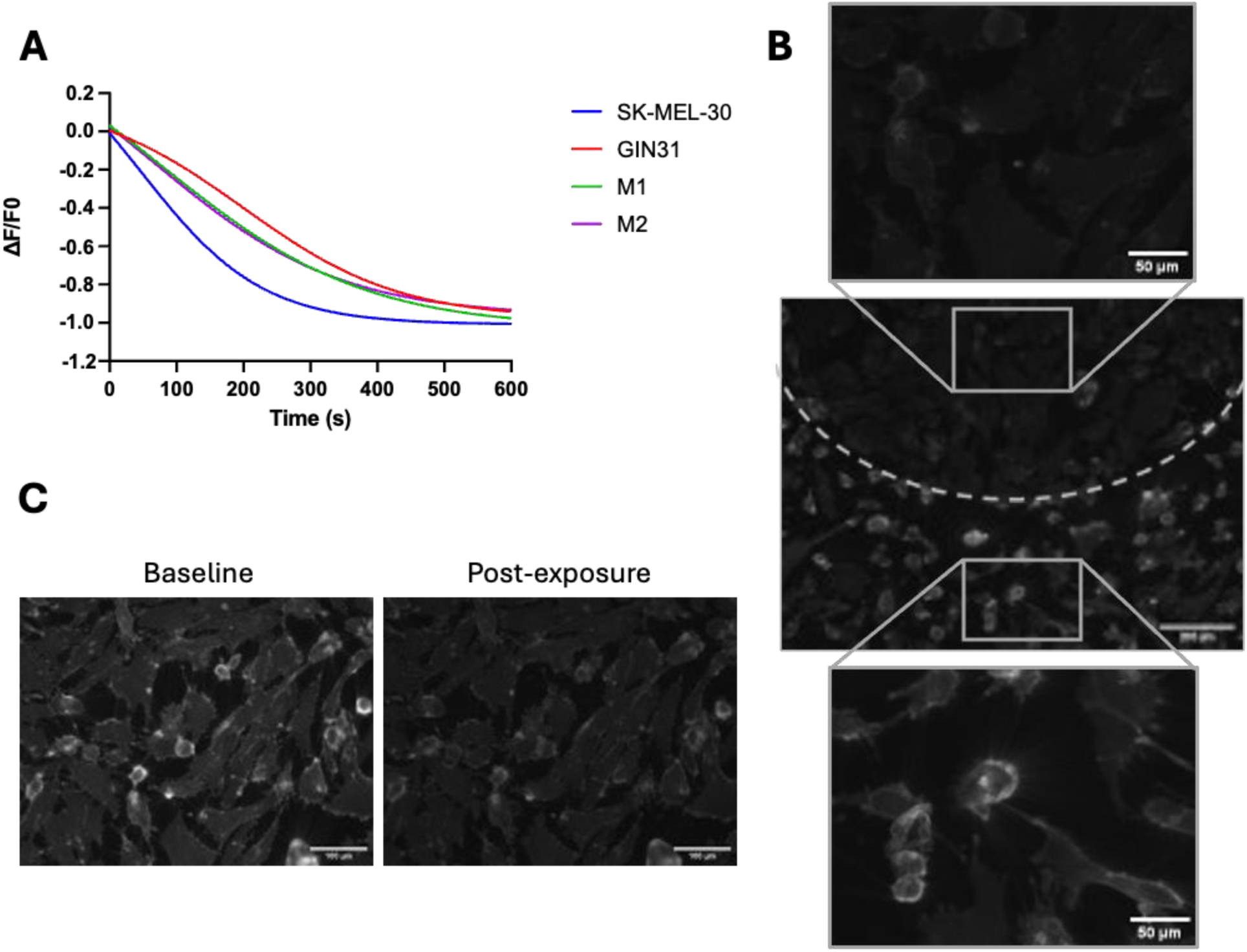
FluoVolt photobleaching varies by cell type. **A.** Cells were imaged on a Nikon Eclipse Ti2 every 10 ms for 10 minutes. Fluorescence intensity was quantified and expressed as ΔF/F₀, where F₀ represents the initial fluorescence intensity. Data represent the mean of three independent biological replicates. Solid lines represent nonlinear regression fits to a four-parameter logistic (4PL) model, demonstrating varied rates of photobleaching between cell types. **B.** Representative images of GIN31 cells following exposure. In the central image, the dotted line indicates the extent of 488 laser exposure, with the upper and lower panels showing magnified views of exposed and non-exposed regions, respectively. **C.** Representative images of GIN31 cells at baseline and 10 minutes after 488 laser exposure.

For experiments involving more than two groups (Figures 2 and 3), statistical significance was determined using one-way analysis of variance (ANOVA) with Šídák’s correction for multiple comparisons for cancer cell lines. For Mφ experiments, repeated-measures one-way ANOVA was used to account for donor-dependent variability.

**Figure 2:**
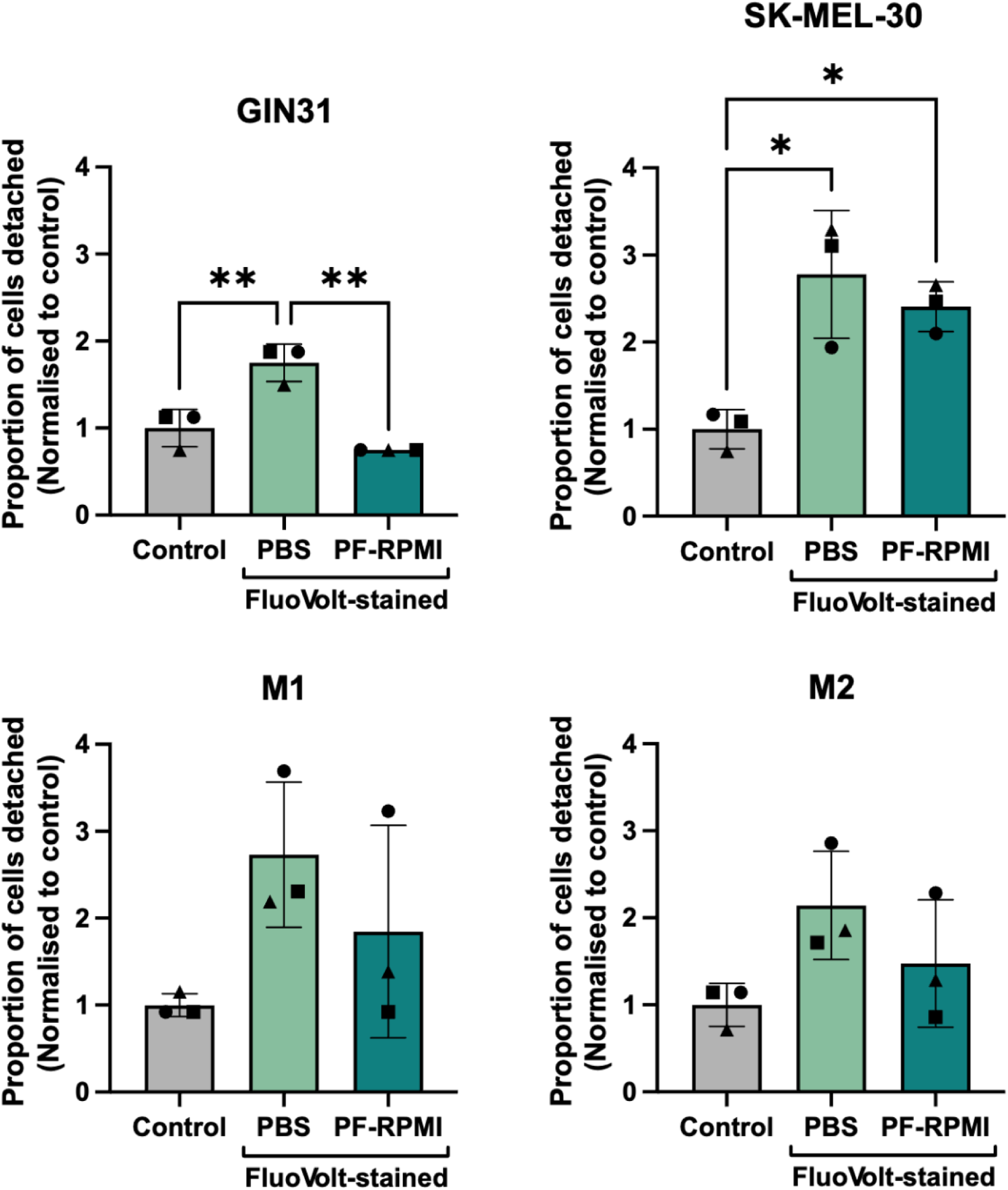
FluoVolt-induced cell detachment is reduced in phenol red-free medium. GIN31 and SK-MEL-30 cells and hMDMs were either stained with FluoVolt or incubated in PBS as a control and imaged under laser exposure in either PBS or phenol red-free medium (PF-RPMI). In GIN31 and SK-MEL-30 cells, FluoVolt staining increased cell detachment in PBS compared to control conditions, which was reduced in PF-RPMI. A similar trend was observed in hMDMs but did not reach statistical significance. Bars represent mean ± SEM, with individual symbols indicating biological replicates or donors. Statistical significance was determined using one-way ANOVA (cancer cell lines) or repeated-measures one-way ANOVA (hMDMs) with Šídák’s multiple comparisons test (* = *p* ≤ 0.05; ** = *p* ≤ 0.01; non-significant comparisons not shown).

**Figure 3:**
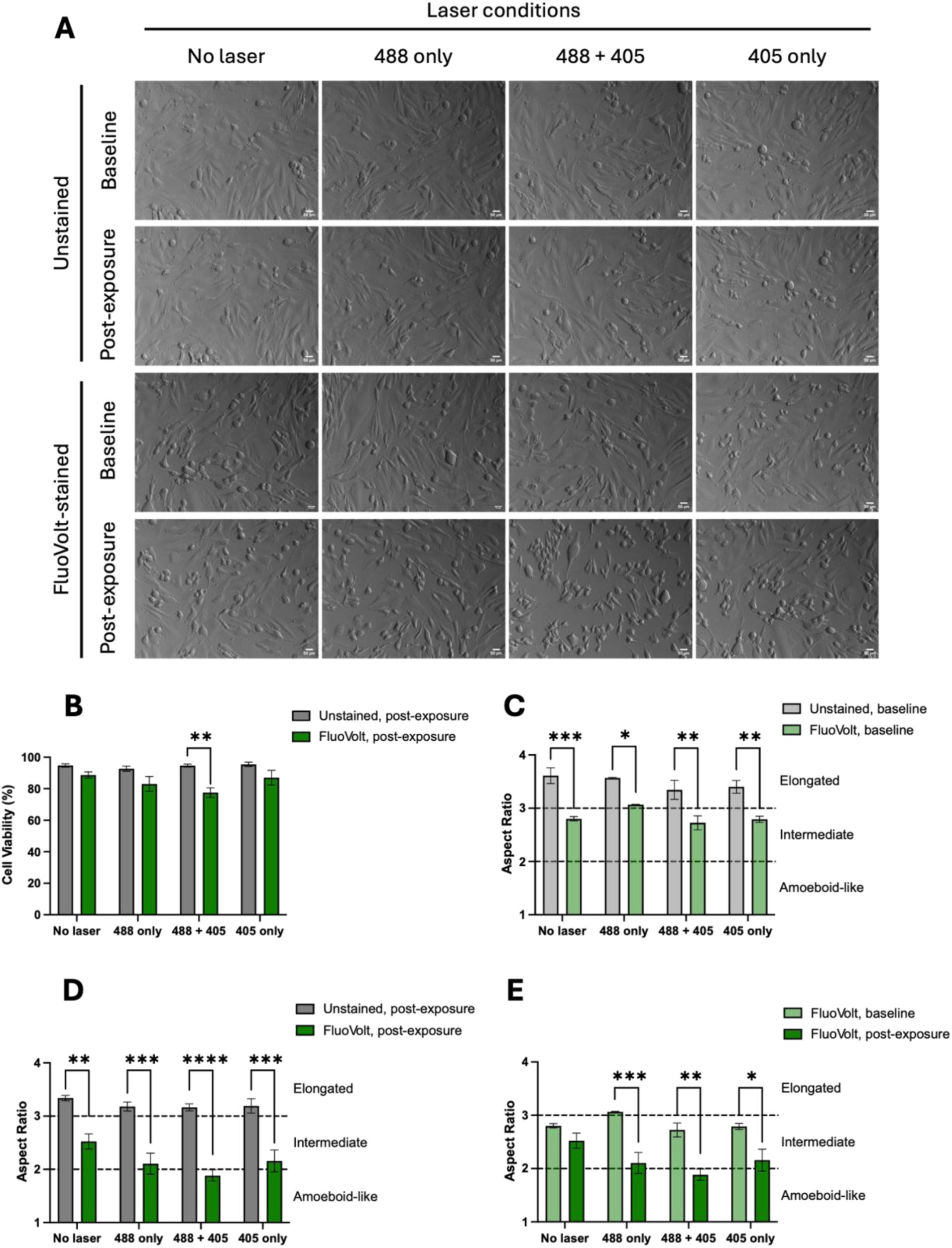
FluoVolt reduces cell viability and alters cell morphology. **A.** Representative images of GIN31 cells, either unstained or stained with FluoVolt and imaged at initial laser exposure (baseline) and after 10 minutes of rest (post-exposure). Cells were imaged under four conditions: no laser exposure, 488 laser only, 488 and 405 lasers, and 405 laser only. **B.** FluoVolt-stained GIN31 cells show reduced viability upon combined 488 and 405 laser exposure. **C.** Across all conditions, FluoVolt staining reduced GIN31-cell aspect ratio even in the absence of laser exposure, indicating a shift from an elongated to an intermediate morphology. **D.** In FluoVolt-stained GIN31 cells, laser exposure further altered aspect ratio compared to unstained cells post-exposure, with dual laser excitation promoting an ameboid-like phenotype. **E.** Laser exposure amplified FluoVolt-induced morphological changes compared with unstimulated GIN31 cells. **For all graphs:** Bars represent mean ± SEM across 3 biological repeats. One-way ANOVA with Šídák’s correction for multiple comparisons (* = *p* ≤ 0.05; ** = *p* ≤ 0.01; *** = *p* ≤ 0.001; **** = *p* ≤ 0.0001; non-significant comparisons not shown).

For comparisons between two groups (Figure 4), statistical significance was assessed using Welch’s unpaired t-test to account for unequal variances between conditions.

**Figure 4:**
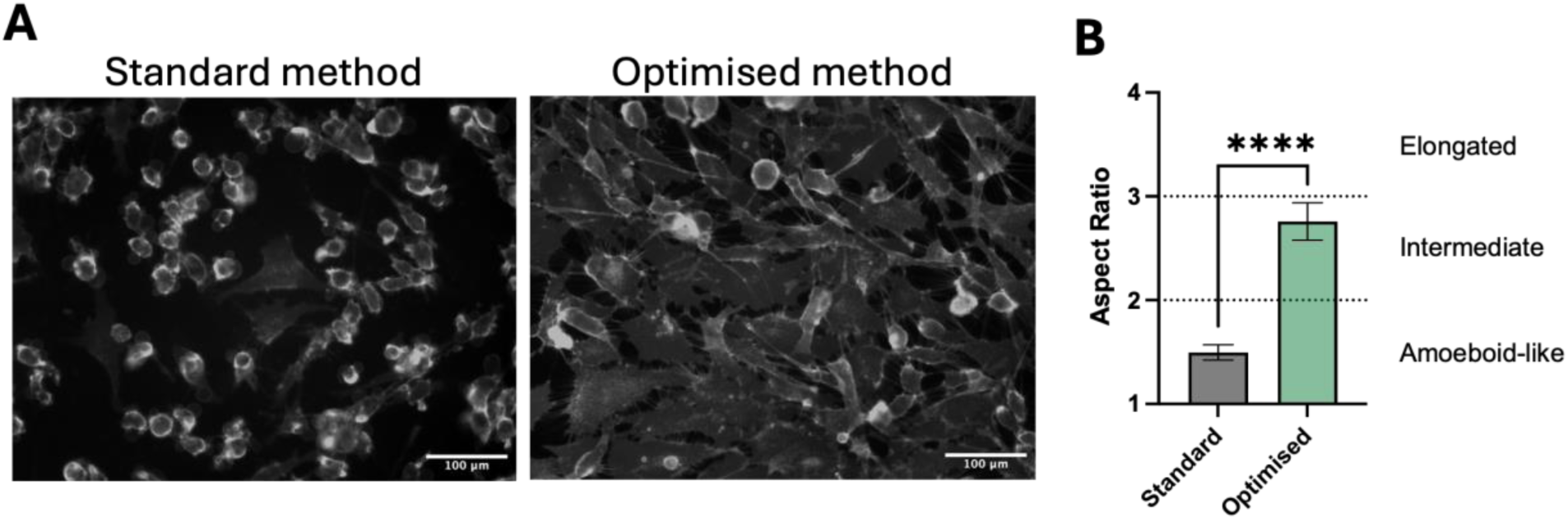
Reduced FluoVolt concentration and incubation time mitigate effects on cell morphology. **A.** Representative images of GIN31 cells stained using reduced concentration and incubation time (half concentration, half incubation) compared to the standard method, 10 minutes after initial laser exposure, show decreased morphological alterations. **B.** Quantification by aspect ratio analysis confirmed a reduction in these effects under modified conditions. Bars represent mean ± SEM from 25 cells analysed from a single biological repeat. Statistical significance was determined using Welch’s unpaired t-test (**** = *p* ≤ 0.0001).

Data are presented as mean ± standard error of the mean (SEM) unless otherwise stated. A *p*-value < 0.05 was considered statistically significant.

## Results

### FluoVolt photobleaching varies by cell type

Initially, we wanted to optimise a protocol for short-term time-course imaging in these cell types. GIN31 and SK-MEL-30 cells and hMDMs, M1 and M2, were stained with FluoVolt and imaged at 10 ms intervals for 10 minutes. Distinct differences in dye-photobleaching kinetics were observed between cell types (Figure 1A), with SK-MEL-30 cells exhibiting complete signal loss within 400 seconds, whereas other cell types retained fluorescence until later time points. Localised laser exposure confirmed that photobleaching was restricted to directly illuminated cells (Figure 1B, upper panel) while cells not directly illuminated maintained their fluorescence intensity (Figure 1B, lower panel). A marked reduction in fluorescence intensity following imaging was evident across all cell types, as shown in Figure 1A, with representative images of GIN31 cells after initial laser exposure (baseline) and 10 minutes post-imaging (post-exposure) shown in Figure 1C.

Cell type-dependent differences in photobleaching, together with sensitivity to imaging conditions, emphasise the importance of optimising experimental parameters for reliable FluoVolt-based measurements.

### FluoVolt staining causes cell detachment

Photobleaching is frequently associated with phototoxicity due to excitation-induced reactive oxygen species generation, leading to altered cellular behaviour, loss of adhesion, morphological changes, and eventual cell death^27–29^.

Given the observed changes in fluorescence stability during imaging, we next sought to determine whether FluoVolt staining and imaging conditions influenced cell adhesion. To investigate this, we assessed the extent of cell detachment following FluoVolt staining. We hypothesised that the use of PBS during imaging may contribute to increased detachment due to its limited capacity to support adhesion compared to complete culture medium. We therefore compared imaging in PBS with phenol red-free medium (PF-RPMI).

GIN31, SK-MEL-30, and hMDMs were either stained with FluoVolt or incubated in PBS as a control, with all steps including washes maintained. FluoVolt-stained cells were then imaged under laser exposure in either PBS or PF-RPMI, while unstained cells were imaged in PBS. Following imaging, the supernatant was collected and detached cells were quantified.

Across all cell types, FluoVolt staining in PBS resulted in an approximately 2.5-fold increase in cell detachment compared to unstained conditions. This increase was statistically significant in both GIN31 (*p* < 0.01) and SK-MEL-30 cells (*p* < 0.05). The use of PF-RPMI mitigated this effect, completely preventing the increase in detachment in GIN31 cells (*p* < 0.01 compared to PBS conditions) and partially reducing it in SK-MEL-30 cells. A similar trend was observed in hMDMs; however, this was not statistically significant, likely due to donor-dependent variability.

These findings identify imaging medium as a critical determinant of cell adhesion during FluoVolt-based experiments and highlight the importance of optimising experimental conditions to minimise cell loss, resulting in artefactual decreases in fluorescence.

### FluoVolt staining and excitation reduces cell viability and alters cell morphology

As imaging medium was found to significantly affect cell adhesion, all subsequent experiment were imaged in PF-RPMI. We next aimed to characterise the broader impact of FluoVolt staining and laser exposure on cell morphology and viability, focusing on GIN31 cells to optimise staining protocols in this cell line.

FluoVolt-stained GIN31 cells exhibited morphological changes that were further exacerbated by laser exposure (Figure 3A). Cells were stained and imaged under brightfield (baseline) and excited with the following: no laser exposure, 488 laser only, 405 laser only, or dual exposure (488+405). Cells were then rested for 10 minutes, and subsequently reimaged under brightfield (post-exposure), revealing pronounced structural alterations. The imaging solution was collected, and the remaining cells were enzymatically dissociated using trypsin. These were then combined and viability quantified using Trypan Blue exclusion, revealing a clear reduction in cell viability following laser exposure.

As shown in Figure 3B, dual excitation with both 488 and 405 lasers reduced cell viability by approximately 17.5% in FluoVolt-stained cells compared to unstained controls (*p* ≤ 0.01), whereas no laser exposure or single-channel excitation did not result in a significant reduction.

Regardless of laser excitation, FluoVolt staining reduced the aspect ratio of cells. To measure their aspect ratio, a minimum of 25 cells per condition were analysed across 3 independent images, with the same 3 fields of view maintained across wells to account for intra-well variability. An aspect ratio > 3 is defined as elongated, values < 2 are defined as amoeboid-like, and values between 2 and 3 defined as intermediate.

Across all conditions, FluoVolt staining reduced the aspect ratio of cells at baseline, shifting them from an elongated phenotype towards an intermediate phenotype (Figure 3C). This effect was further enhanced following laser exposure, with cell morphology progressing towards an amoeboid-like phenotype, particularly under dual excitation (Figure 3D).

In the absence of laser exposure, FluoVolt-stained cells maintained an intermediate phenotype over time, whereas laser excitation promoted a transition towards an ameboid-like phenotype, with dual excitation producing the most pronounced effect (Figure 3E).

The combined effects of FluoVolt staining and laser exposure drive substantial alterations in cell morphology and viability, underscoring the importance of optimising experimental parameters to avoid artefactual phenotypic changes.

### Reducing FluoVolt concentration and incubation time mitigates morphological effects

To address the effects of FluoVolt staining and imaging conditions on cell morphology and viability, we next investigated whether these perturbations could be minimised through optimisation of staining parameters.

Cells were stained using half of the standard dye concentration (1:2000) and PowerLoad concentration (1:200) and a shortened incubation time (10 minutes versus 20 minutes), to mitigate potential phototoxic effects, and imaged in PF-RPMI, to minimise cell detachment. Following laser exposure and a 10-minute recovery period, cells were imaged and analysed for morphological changes using aspect ratio measurements (Figure 4).

Under standard staining conditions, cells predominantly exhibited an amoeboid-like phenotype, characterised by lower aspect ratios. In contrast, cells stained under reduced conditions retained a more intermediate phenotype, indicating diminished morphological disruption. This was confirmed by aspect ratio analysis, which demonstrated a significant attenuation of morphological effects under modified conditions compared to standard staining (*p* < 0.0001).

These findings establish modified staining conditions as an effective approach to reduce FluoVolt-induced artefacts while preserving physiologically relevant cell morphology.

## Discussion

Fluorescent VSDs such as FluoVolt provide a powerful alternative to electrophysiological techniques, enabling non-invasive and high-throughput measurements of V_mem_ across large cell populations^17,18^. However, the findings of this study demonstrate that FluoVolt staining and imaging can introduce significant artefacts, including photobleaching, cell detachment, reduced viability, and pronounced morphological changes. These effects have important implications for the interpretation of fluorescence-based V_mem_ measurements, particularly under prolonged imaging conditions.

Photobleaching was observed in all cell types but occurred at markedly different rates, with SK-MEL-30 cells exhibiting rapid fluorescence decay throughout imaging compared to GIN31 cells and hMDMs. This suggests that photobleaching is not solely a property of the dye itself but is strongly influenced by cellular context. Differences in membrane composition, dye uptake, and intracellular environment may all contribute to the observed variability^20^. Such cell-type-dependent behaviour highlights the importance of validating dye performance for each experimental system rather than assuming uniform behaviour across models^22^.

In addition to photobleaching, FluoVolt staining in combination with laser exposure resulted in increased cell detachment across all cell types. We postulated that the use of PBS during imaging may have exacerbated this effect, given its reduced capacity to support adhesion relative to complete culture medium. The use of phenol red-free medium in this study was primarily intended to minimise background fluorescence and improve signal detection, as phenol red is known to contribute to increased background signal in fluorescence imaging^30^. Together, these considerations highlight the importance of carefully optimising imaging conditions to both reduce biological artefacts and improve signal-to-noise ratios.

The observed cell detachment is consistent with previous reports in which prolonged excitation of FluoVolt-labelled human induced pluripotent stem cells led to cell retraction, membrane blebbing, and eventual cell death^31^. More broadly, phototoxicity in fluorescence imaging is widely attributed to the generation of reactive oxygen species during excitation, which can induce oxidative damage, disrupt membrane integrity, and impair cell adhesion^28^.

In addition to light-induced effects, the chemical properties of the dye itself may contribute to cellular perturbation. FluoVolt loading relies on AM ester hydrolysis and the use of dispersal agents such as Pluronic F127, which has been reported to generate potentially toxic by-products, including formaldehyde, during intracellular processing^32^. This may introduce baseline cellular stress even in the absence of prolonged illumination, consistent with the morphological changes observed in stained cells without laser exposure.

FluoVolt staining was also associated with reduced cell viability and significant alterations in cell morphology, particularly following laser exposure. Dual laser excitation of FluoVolt-labelled cells reduced cell viability by approximately 20% compared to unstained cells under the same laser excitation and approximately 12% relative to FluoVolt-stained cells without laser excitation, supporting a light dose-dependent mechanism of phototoxicity. Such dose dependence is well-established in fluorescence imaging, where increased excitation intensity and duration amplify phototoxic damage^33,34^. In terms of morphology, cells transitioned from elongated to intermediate or amoeboid-like phenotypes, indicative of cytoskeletal remodelling and cellular stress responses^28^. Notably, these changes were evident even in the absence of laser exposure, suggesting that dye loading alone can perturb cellular behaviour, with subsequent illumination exacerbating these effects.

To mitigate these issues, we evaluated modified staining conditions using reduced dye and loading solution concentration, and reduced incubation time. These optimised conditions significantly decreased the impact of FluoVolt on cell morphology. Although this approach was selected based on direct comparison of representative conditions rather than exhaustive optimisation, the results demonstrate that relatively simple adjustments can substantially reduce dye-induced artefacts. This highlights the importance of empirically optimising staining protocols for each experimental system, particularly when studying processes sensitive to changes in cell morphology or viability.

The extent of photobleaching and phototoxicity observed is also likely to depend on the imaging modality employed. Previous studies have shown that two-photon excitation of FluoVolt reduces both photobleaching and phototoxic effects compared to conventional widefield or confocal microscopy, emphasising the role of excitation strategy in determining experimental outcomes^35^. Together with our findings, this suggests that both staining conditions and imaging parameters must be carefully considered to minimise perturbation.

Collectively, this study demonstrates that, while FluoVolt is a valuable tool for optical measurement of V_mem_, its use is associated with significant context-dependent effects on cell behaviour. Without appropriate optimisation, these effects may confound experimental readouts, particularly in long-term imaging experiments. The dependence of these artefacts on cell type, imaging environment, dye concentration, and excitation conditions underscores the need for careful validation in each experimental context and use of appropriate controls. By identifying key sources of variability and demonstrating practical strategies to mitigate them, this work provides guidance for the more reliable application of FluoVolt and related VSDs in live-cell imaging.

## Conclusions

In this study, we demonstrate that FluoVolt staining and fluorescence imaging can significantly influence cell behaviour, producing photobleaching, reduced viability, increased detachment, and pronounced morphological changes. These effects were strongly dependent on cell type, imaging conditions, and excitation parameters, highlighting the context-specific nature of VSD performance. Importantly, optimisation of staining conditions through reduced dye concentration and incubation time substantially mitigated these effects, while maintaining measurable fluorescence signals. Our findings emphasise that, although FluoVolt is a powerful tool for optical membrane V_mem_ measurements, its application requires careful experimental validation to avoid confounding artefacts. This work provides practical guidance for minimising dye-induced perturbation and supports the more reliable use of VSDs in live-cell imaging applications.

## AUTHOR INFORMATION

### Author Contributions

The manuscript was written through contributions of all authors. All authors have given approval to the final version of the manuscript. Writing, E.M.A., M.I.M., F.G., F.P., R.R., S.J.S., I.S., J.M.R., H.F., A.M.J., A.M.B., A.A.M. and F.J.R. Data Curation, Investigation, and Analysis E.M.A., M.I.M., F.G., F.P. Funding acquisition, A.A.M and F.J.R. Supervision, A.A.M., F.J.R., and A.M.B.

### Data availability

All other data associated with this manuscript can be found at DOI: 10.17639/nott.26020.

## ABBREVIATIONS

4PL: four-parameter logistic
5-ALA: 5-aminolevulinic acid
ANOVA: analysis of variance
DMEM: Dulbecco’s modified Eagle medium
DPBS: Dulbecco’s phosphate-buffered saline
EDTA: ethylenediaminetetraacetic acid
FBS: foetal bovine serum
FITC: fluorescein isothiocyanate
GM-CSF: granulocyte-macrophage colony-stimulating factor
hMDMs: human monocyte-derived macrophages
M-CSF: macrophage colony-stimulating factor
Mφ: macrophage
PBS: phosphate-buffered saline
PET: photoinduced electron transfer
PF-RPMI: phenol-red free Roswell Park Memorial Institute 1640 medium
RPMI: Roswell Park Memorial Institute 1640 medium
SEM: standard error of the mean
V_mem_: membrane potential
VSD: voltage-sensitive dye.

